# Temporal and spatial limitations in global surveillance for bat filoviruses and henipaviruses

**DOI:** 10.1101/674655

**Authors:** Daniel J. Becker, Daniel E. Crowley, Alex D. Washburne, Raina K. Plowright

## Abstract

Sampling reservoir hosts over time and space is critical to detect epizootics, predict spillover, and design interventions. However, because sampling is logistically difficult and expensive, researchers rarely perform spatiotemporal sampling of many reservoir hosts. Bats are reservoirs of many virulent zoonotic pathogens such as filoviruses and henipaviruses, yet the highly mobile nature of these animals has limited optimal sampling of bat populations. To quantify the frequency of temporal sampling and to characterize the geographic scope of bat virus research, we here collated data on filovirus and henipavirus prevalence and seroprevalence in wild bats. We used a phylogenetically controlled meta-analysis to next assess temporal and spatial variation in bat virus detection estimates. Our analysis shows that only one in four bat virus studies report data longitudinally, that sampling efforts cluster geographically (e.g., filovirus data are available across much of Africa and Asia but are absent from Latin America and Oceania), and that sampling designs and reporting practices may affect some viral detection estimates (e.g., filovirus seroprevalence). Within the limited number of longitudinal bat virus studies, we observed high heterogeneity in viral detection estimates that in turn reflected both spatial and temporal variation. This suggests that spatiotemporal sampling designs are essential to understand how zoonotic viruses are maintained and spread within and across wild bat populations, which in turn could help predict and preempt risks of zoonotic viral spillover.

## Introduction

Risks of pathogen spillover vary across time and space [1,2], in part because pathogen shedding from reservoir hosts is a dynamic spatiotemporal processes [3,4]. Metapopulation dynamics and other spatial processes characterize many reservoir hosts [5], where populations connectivity can determine the spatiotemporal distribution of a pathogen [6,7] and degree of spatial synchrony structuring infection dynamics [8]. Temporal pulses of shedding driven by seasonality in birth and climate are also common [9,10]. Understanding how infection in reservoir hosts varies over space and time is thus a critical need for predicting and managing zoonotic disease risks.

However, surveillance strategies often do not sample this underlying spatiotemporal process, as spatially and temporally explicit designs present logistical challenges when studying mobile and gregarious species [3,11,12]. For hosts such as birds and bats, surveillance is often opportunistic or relies on convenience sampling [13]. These non-probabilistic and often single sampling events cannot characterize spatial and temporal fluctuations in infection, can over- or under-represent times or locations of high prevalence, and can result in non-randomly missing data [3,14]. These challenges cannot be fixed with statistical modeling and can bias estimates of prevalence and epidemiological parameters such as the basic reproductive number [13,15].

Given a fixed cost, difficult decisions must be made about how to allocate sampling efforts. Sampling over space facilitates detecting geographic clusters of disease and predictive mapping [16,17], while sampling over time can identify periods of intensive pathogen shedding and enable inference about dominant transmission routes [18,19]. Researchers often treat this as a tradeoff between sampling over either time or space, rather than allocating effort to both [20]. Implicit here is that the temporal component is constant over space or that the spatial component is constant over time, and such sampling designs result in no data to assess this assumption.

We here quantify the temporal and spatial data limitations for two taxa of high-profile zoonotic viruses of bats: the family *Filoviridae* and genus *Henipavirus*. Bats have been widely studied as reservoirs for zoonotic pathogens and host more viruses with zoonotic potential than other mammals [21,22]. Henipaviruses and some filoviruses (e.g., Marburg virus) can be shed from bats into the environment [23,24] and can cause fatal disease in humans by environmental exposure or from contact with intermediate hosts such as horses, wild primates, or pigs [25–30]. Current evidence suggests many filo- and henipaviruses show variable dynamics in space and time, including shedding pulses from bats [6,25,31–33], which implies that spatiotemporal sampling is critical to capture viral dynamics in bat reservoirs. Yet while past efforts have focused on bat virus discovery [34], the determinants of reservoir status [35], and experimental mechanisms of viral transmission [36], spatiotemporal studies of bat–virus dynamics are rare [37]. This limits understanding how zoonotic viruses are maintained and spread within and across bat populations and impairs improving future sampling designs and ecological interventions [20,38]. We here systematically collated data on the prevalence and seroprevalence of filo- and henipaviruses in wild bats to (*i*) quantify the frequency of temporal studies and (*ii*) assess the geographic scope of current research. We used phylogenetic meta-analysis to (*iii*) quantify how sampling designs and reporting practices may influence viral detection estimates. Single snapshots could miss pulses of viral shedding from bats, whereas pooling data over time could under- or overestimate viral presence [18,20]. Lastly, we (*iv*) characterized the degree of temporal and spatial variation in bat virus detection estimates.

## Methods

To systematically identify studies quantifying the proportion of wild bats positive for filoviruses and henipaviruses using PCR or serology, we searched Web of Science, CAB Abstracts, and PubMed (see Fig. S1). Our dataset included 1176 records from 68 studies. Viruses included not only Hendra, Nipah, Ebola, and Marburg virus but also Lloviu and Reston virus. We grouped viruses by taxa given our sample sizes and known issues of serological cross-reactivity [39,40].

From each study, we defined sampling subunits: a temporally defined sampling event of one bat species in one location per viral detection estimate. Each subunit is the lowest spatial, temporal, and phylogenetic scale (of bats and their viruses) reported. We classified subunits into three sampling designs and reporting practices: one sampling event, multiple events, or pooled events over time. Records of a single prevalence or seroprevalence estimate from a population sampled from a period less than or equal to one month were classified as single sampling events, whereas records of a population over multiple monthly timepoints were classified as spanning multiple events (i.e., a longitudinal study). For example, every monthly prevalence estimate per population of *Pteropus lylei* in Thailand would represent a unique subunit and be classified as longitudinal [41]. Records of a period longer than one month were classified as pooled events, where researchers may have sampled a population across more than one timepoint but reported data as a single viral detection estimate. A schematic of these categorizations is provided in Figure 1A. One month was selected because this timeframe was the lowest common temporal unit and because bat shedding of these viruses can occur within a month [36,42]. These data were reported for most records (1121/1176 subunits; three publications did not report these data and three additional publications did not always report such data for all records). For each subunit, we also recorded the bat species, virus taxon, coarse detection method (i.e., PCR or serology), number of bats sampled, proportion of bats positive, sampling timepoints, sampling location, and country (recoded to the United Nations geoscheme for our descriptive analyses).

**Figure 1.**
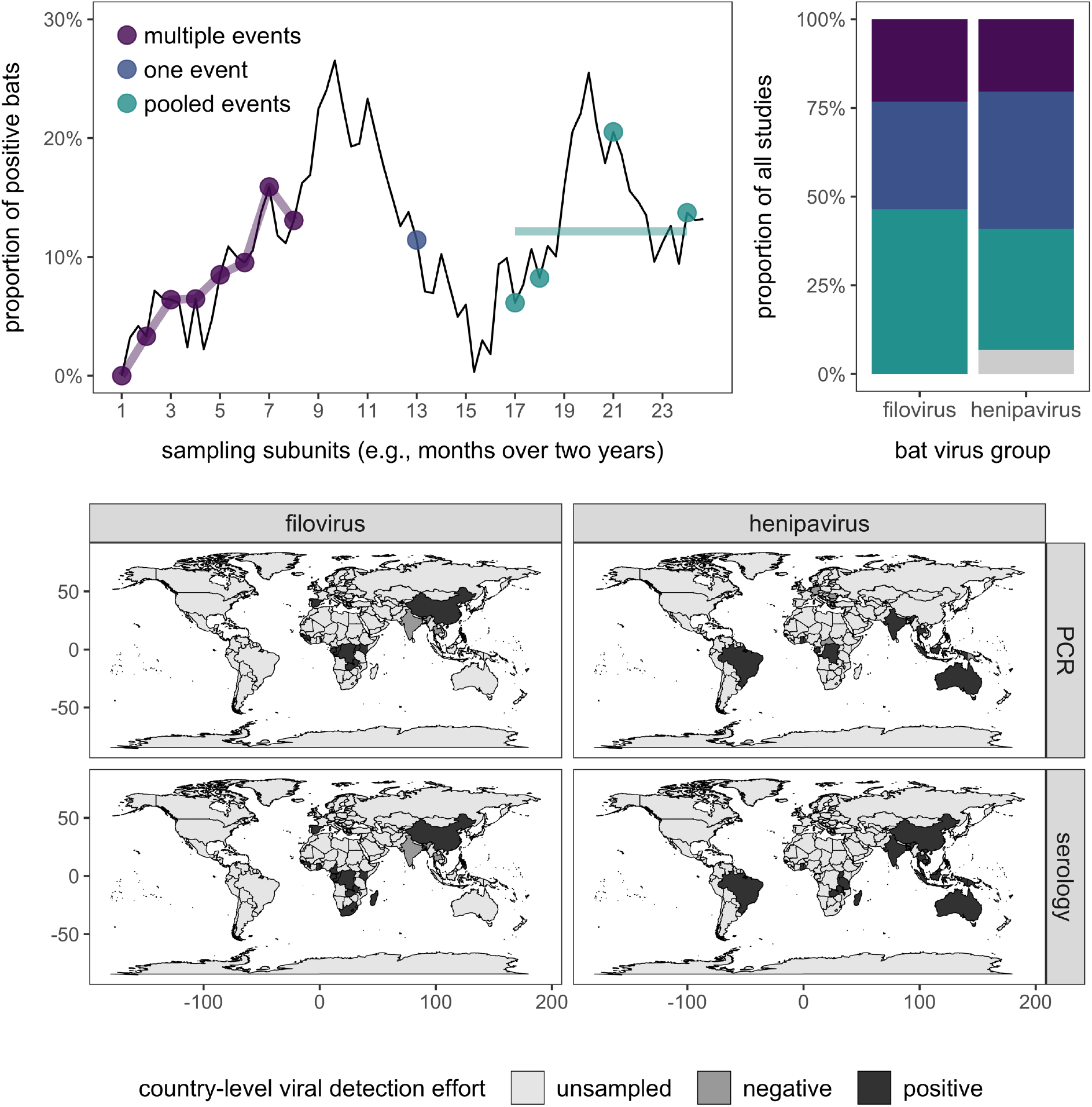
Top: Conceptual schematic of how different sampling designs and reporting practices (colored points and lines) capture the underlying temporal dynamics of infection (black line), followed by observed proportions for studies of bat filoviruses and henipaviruses (grey shows the proportion of studies not reporting these data). Bottom: Countries sampled for bat filoviruses and henipaviruses and where wild bats have been found positive through PCR or serology.

We quantified the proportion of studies using each sampling and reporting design, both across all data and stratified by virus taxon. To assess how the frequency of longitudinal studies (i.e., those with repeated sampling) has changed over time, we fit a generalized additive model with the *mgcv* package in R and a smooth term for publication year [43]. We also calculated the duration of repeat sampling for these longitudinal studies. For studies that pooled data over time, we quantified days represented per subunit. To describe geographic biases in bat virus studies, we assessed sampling gaps according to region (United Nations geoscheme). We used a χ^2^ test to assess if sampling designs and reporting practices were differently distributed across regions.

To assess the contribution of sampling designs and reporting practices to viral detection estimates and to quantify the degree of spatial and temporal variation in bat–virus interactions, we used the *metafor* package to calculate logit-transformed proportions and sampling variances and to fit hierarchical meta-analysis models [44,45]. To account for phylogenetic dependence, we included bat species as a random effect [46], for which the covariance structure used the phylogenetic correlation matrix; we obtained our phylogeny from the Open Tree of Life with the *rotl* and *ape* packages [47,48]. We excluded subunits that pooled data across or within bat genera (*n*=102). As few subunits (*n*=14) pooled data across specified species in a genus, we randomly selected one species to retain these records. Our final dataset included 1019 subunits from 60 studies and 215 bat species (Fig. S2). Our models also included subunit nested within study as a random effect and weighting by sampling variances. To first assess heterogeneity among viral detection estimates, we fit a random-effects model (REM; intercept only) and stratified this analysis per viral taxon and detection method. We used restricted maximum likelihood to obtain unbiased estimates of the variance components, from which we derived *I*^*2*^ to quantify the contribution of true heterogeneity to total variance in viral detection estimates [49]. We used these estimates to partition variance attributed to each random effect; in the case of bat species, we derived phylogenetic heritability (*H*^*2*^) as a measure of phylogenetic signal [46]. We used Cochran’s *Q* to test if such heterogeneity was greater than expected by sampling error alone [50].

To next test how sampling designs and reporting practices may influence viral detection estimates, we fit a mixed-effects model (MEM) with the same random effects and an interaction between sampling design and reporting practices, detection method, and virus taxon. We tested significance of moderators and interactions using the *Q* test [44] and derived a pseudo-*R*^*2*^ as the proportional reduction in the summed variance components compared with those of a REM [51].

To test if viral detection estimates showed spatiotemporal variation, we fit models with the same random effects to our data subset reporting multiple events (*n*=273). We fit a REM to quantify *I*^*2*^ for longitudinal studies. We then fit MEMs with location and month as univariate moderators to test if viral detection estimates varied across space and time. Because this subset of the data included many unique locations (*n*=28) and months (*n*=12), we did not use interaction terms and instead fit an additional set of MEMs to each viral taxon–detection method strata.

## Results

Only 26% of bat virus studies reported data longitudinally (10 filo- and 9 henipavirus studies; Fig. 1). However, the frequency of such studies has weakly increased over time (Fig. S3, χ^2^_1_=2.75, *p*=0.1). Eleven studies reported sampling populations 2–3 times while 12 reported sampling populations over four times. The duration of longitudinal studies ranged from 150 days to over 10 years, on average spanning 2.5 years of repeat sampling (Fig. S4). In contrast, half of our studies (*n*=34) instead reported estimates across multiple timepoints as pooled proportions, which on average represented 644 days of temporally aggregated data (SD=492; Fig. S5).

Bat sampling also showed geographic biases (Fig. 1, Table 1). Filovirus studies were conducted across much of Africa and Asia but not in Latin America and Oceania. PCR and serology have been used in the same region in most areas, but only one or the other have been used in Southern Africa for filoviruses and in Europe, Eastern and Middle Africa, and Eastern Asia for henipaviruses (Table 1). Geography was also associated with sampling design and reporting practices (*χ*^*2*^=369.3, *p*=0.001). Longitudinal data were only reported from Central, Eastern, Middle, and Southern Africa for filoviruses and only reported from Southeastern Asia, Eastern Africa, and Oceania for henipaviruses (Table 1).

**Table 1.** Summary of the temporal and spatial limitations for bat filovirus and henipavirus prevalence and seroprevalence data. Some studies had multiple diagnostic methods, sampling designs, and reporting methods. Diagnostic mismatch refers to geographic regions (United Nations geoscheme) where either PCR or serology have been used (but not together).

|  |  | Longitudinal virus studies | Geographic sampling gaps | Diagnostic mismatch | Regions with longitudinal data |
| --- | --- | --- | --- | --- | --- |
| Filoviruses | PCR | 4/19 | Latin America, Oceania, Southern Africa | Southern Africa | Central Africa, Eastern Africa |
|  | Serology | 7/25 | Latin America, Oceania |  | Central Africa, Eastern Africa, Middle Africa, Southern Africa |
| Henipaviruses | PCR | 4/13 | Eastern Africa, Southern Africa, Eastern Asia | Europe, Eastern Africa, Middle Africa, Eastern Asia | Southeastern Asia, Oceania |
|  | Serology | 5/27 | Middle Africa, Southern Africa, Europe |  | Southeastern Asia, Oceania, Eastern Africa |

We observed significant heterogeneity across viral detection estimates (*I*^*2*^=0.91, *Q*_*1017*_=6929, *p*<0.001). Bat species and study accounted for most variation (*I*^*2*^_*species*_=0.41, *I*^*2*^_*study*_=0.34, *H*^*2*^=0.45; Table S2). We also found significant heterogeneity within each viral taxon–detection strata, although *I*^*2*^ and *H*^*2*^ values varied across these subsets (Table S1). Viral detection estimates for henipaviruses had much stronger phylogenetic signal than filoviruses.

Our MEM showed that viral detection estimates broadly varied with detection method (*Q*_*1*_=5.41, *p*=0.02; seroprevalence was generally higher than prevalence) and were associated with sampling design and reporting; however, the effect tended to depended on virus taxa and detection method (three-way interaction: *Q*_*2*_=5.36, *p*=0.07, *R*_*2*_=0.06; Table S2). A post-hoc analysis with MEMs fit to each strata showed sampling design and reporting were associated with filovirus seroprevalence (*Q*_*2*_=10.30, *p*=0.006; Fig. 2), with longitudinal studies generally showing higher proportions of positive bats. Sampling design and reporting had no effects on henipavirus seroprevalence nor prevalence estimates for either virus taxon (Table S3).

**Figure 2.**
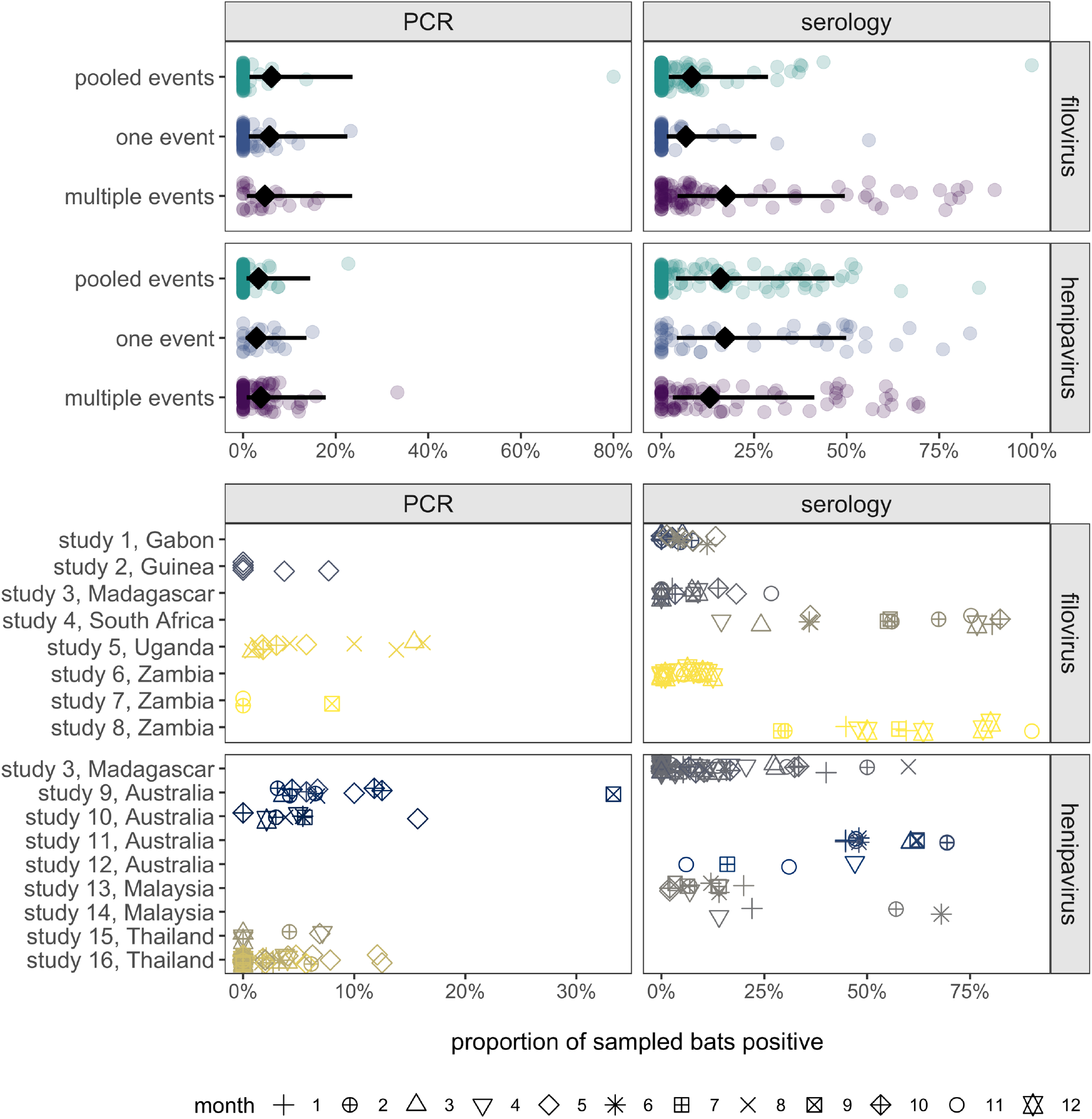
Top: Influence of sampling designs and reporting practices on virus detection estimates. Points show proportions of positive bats per subunit; lines and diamonds display back-transformed predicted means and 95% confidence intervals from the MEM. Bottom: Spatiotemporal variation in viral detection estimates for longitudinal studies. Points represent each subunit virus detection estimate and are colored by locations and shaped by month.

We also detected high variation in viral detection estimates across longitudinal studies (*Q*_*271*_=2866, *p*<0.0001, *I*_*2*_=0.94; Fig. 2). Study contributed more to residual variance than phylogeny (*I*^*2*^_*species*_=0.28, *I*^*2*^_*study*_=0.52, *I*^*2*^_*subunit*_=0.13). Across these data, location did not predict viral detection estimates (*Q*_*28*_=17.67, *p*=0.91); however, MEMs fit to each strata showed high spatial variation for all subsets except filovirus prevalence (Table S4). Month also had little predictive power across all longitudinal data (*Q*_*11*_=6.93, *p*=0.80), but separate MEMs revealed high temporal variation for filovirus seroprevalence and henipavirus prevalence (Table S5).

## Discussion

Our study provides a systematic synthesis of prevalence and seroprevalence for bat filoviruses and henipaviruses that can guide future sampling. Only one in four studies reported longitudinal data, although use of such approaches is increasing. Half of studies instead pooled data over time (and space). Geographic limitations were also evident, especially for where longitudinal studies have been conducted. This was especially evident for filoviruses; although the absence of studies in Latin America and Oceania may reflect the lack of reported human cases, bat reservoirs are predicted to occur in both regions [35]. Many studies also used either PCR or serology, although using both may improve statistical inference about how zoonotic pathogens persist in hosts [18].

We found generally weak evidence that such variation in sampling design and reporting affected viral detection estimates, although filovirus seroprevalence tended to be greatest from longitudinal studies. Serological surveys of Marburg and Ebola virus have found strong temporal dynamics that may reflect seasonality in bat reproduction or resource availability [31,52,53]. Detection estimates could be higher with repeated sampling, given that such studies are more likely to detect shedding pulses and pooling of data could increase zeros in the numerator (underestimating seroprevalence). The lack of a similar pattern for filovirus PCR data could result from low prevalence and be biased by zero inflation. We also qualify that our low *R*^*2*^, alongside high contributions of bat phylogeny and study random effects, suggests other aspects of bat ecology (e.g., seasonal birth [31,54]) or study idiosyncrasies (e.g., serological cutoffs [39,40]) may play more critical roles in shaping viral detection estimates. High *H*^*2*^ for henipaviruses suggests cladistic or trait-based analyses of shedding could be insightful [35,55]. Yet given the potential for sampling design and reporting to affect viral detection estimates, we encourage researchers to publish data at the lowest spatial, temporal, and phylogenetic scale associated with sampling and to provide data at such scales to facilitate these future analyses.

Our analysis of longitudinal studies found significant spatial and temporal variation in some bat virus data. This implies spatiotemporal sampling is critical to make inferences about bat virus spillover. Although sampling over space and time is challenging, especially for highly mobile animals like bats, sampling can be informed by spatiotemporal variation in prevalence and seroprevalence and analyses of spatiotemporal autocorrelation [20,56]. Greater variation over space can require more fine-scale spatial sampling, and greater variation over time can require more fine-scale temporal sampling. Spatiotemporal designs, such as stratified random sampling or rotating panels, can help capture spatial and temporal variation in virus shedding while also addressing some logistical challenges [13,57,58]. The increased adoption of such approaches, especially in the understudied regions identified in our analysis, will be key to improve understanding bat virus dynamics and how spillover risk varies over time and space.

## Data availability

Data are available in the Dryad Digital Repository [59].

## Author contributions

DJB and DEC designed the study, DEC collected data, DJB analyzed data, and all authors contributed to writing the manuscript.

## Acknowledgements

We thank Megan Higgs and anonymous reviewers for helpful comments on this manuscript.

## References

1. Plowright RK, Parrish CR, McCallum H, Hudson PJ, Ko AI, Graham AL, Lloyd-Smith JO. 2017 Pathways to zoonotic spillover. Nat. Rev. Microbiol. 15, 502–510.

2. Schmidt JP, Park AW, Kramer AM, Han BA, Alexander LW, Drake JM. 2017 Spatiotemporal Fluctuations and Triggers of Ebola Virus Spillover. Emerg. Infect. Dis. 23, 415–422. (doi:10.3201/eid2303.160101)

3. Hoye BJ, Munster VJ, Nishiura H, Klaassen M, Fouchier RAM. 2010 Surveillance of Wild Birds for Avian Influenza Virus. Emerg. Infect. Dis. 16, 1827–1834. (doi:10.3201/eid1612.100589)

4. Plowright RK et al. 2015 Ecological dynamics of emerging bat virus spillover. Proc R Soc B 282, 20142124. (doi:10.1098/rspb.2014.2124)

5. Vergara PM, Saravia-Zepeda A, Castro-Reyes N, Simonetti JA. 2016 Is metapopulation patch occupancy in nature well predicted by the Levins model? Popul. Ecol., 1–9. (doi:10.1007/s10144-016-0550-5)

6. Plowright RK, Foley P, Field HE, Dobson AP, Foley JE, Eby P, Daszak P. 2011 Urban habituation, ecological connectivity and epidemic dampening: the emergence of Hendra virus from flying foxes (Pteropus spp.). Proc. R. Soc. B Biol. Sci. 278, 3703–3712. (doi:10.1098/rspb.2011.0522)

7. Hess G. 1996 Disease in Metapopulation Models: Implications for Conservation. Ecology 77, 1617–1632. (doi:10.2307/2265556)

8. Bjørnstad ON, Ims RA, Lambin X. 1999 Spatial population dynamics: analyzing patterns and processes of population synchrony. Trends Ecol. Evol. 14, 427–432.

9. Altizer S, Dobson A, Hosseini P, Hudson P, Pascual M, Rohani P. 2006 Seasonality and the dynamics of infectious diseases. Ecol. Lett. 9, 467–484.

10. Peel AJ, Pulliam JRC, Luis AD, Plowright RK, O’Shea TJ, Hayman DTS, Wood JLN, Webb CT, Restif O. 2014 The effect of seasonal birth pulses on pathogen persistence in wild mammal populations. Proc. R. Soc. Lond. B Biol. Sci. 281, 20132962. (doi:10.1098/rspb.2013.2962)

11. Kuiken T, Leighton FA, Fouchier R a. M, LeDuc JW, Peiris JSM, Schudel A, Stöhr K, Osterhaus ADME. 2005 Pathogen Surveillance in Animals. Science 309, 1680–1681. (doi:10.1126/science.1113310)

12. Stallknecht DE. 2007 Impediments to Wildlife Disease Surveillance, Research, and Diagnostics. In Wildlife and Emerging Zoonotic Diseases: The Biology, Circumstances and Consequences of Cross-Species Transmission, pp. 445–461. Springer, Berlin, Heidelberg. (doi:10.1007/978-3-540-70962-6_17)

13. Nusser SM, Clark WR, Otis DL, Huang L. 2008 Sampling Considerations for Disease Surveillance in Wildlife Populations. J. Wildl. Manag. 72, 52–60.

14. Venette RC, Moon RD, Hutchison WD. 2002 Strategies and statistics of sampling for rare individuals. Annu. Rev. Entomol. 47, 143–174. (doi:10.1146/annurev.ento.47.091201.145147)

15. Hayman DTS, Suu-Ire R, Breed AC, McEachern JA, Wang L, Wood JLN, Cunningham AA. 2008 Evidence of Henipavirus Infection in West African Fruit Bats. PLOS ONE 3, e2739. (doi:10.1371/journal.pone.0002739)

16. Peterson AT. 2006 Ecologic Niche Modeling and Spatial Patterns of Disease Transmission. Emerg. Infect. Dis. 12, 1822–1826. (doi:10.3201/eid1212.060373)

17. Wang J-F, Stein A, Gao B-B, Ge Y. 2012 A review of spatial sampling. Spat. Stat. 2, 1–14. (doi:10.1016/j.spasta.2012.08.001)

18. Plowright RK, Peel AJ, Streicker DG, Gilbert AT, McCallum H, Wood J, Baker ML, Restif O. 2016 Transmission or Within-Host Dynamics Driving Pulses of Zoonotic Viruses in Reservoir–Host Populations. PLoS Negl. Trop. Dis. 10, e0004796. (doi:10.1371/journal.pntd.0004796)

19. Restif O et al. 2012 Model-guided fieldwork: practical guidelines for multidisciplinary research on wildlife ecological and epidemiological dynamics. Ecol. Lett. 15, 1083–1094.

20. Plowright RK, Becker DJ, McCallum H, Manlove KR. 2019 Sampling to elucidate the dynamics of infections in reservoir hosts. Philos. Trans. R. Soc. B (doi:10.1098/rstb.2018.0336)

21. Luis AD et al. 2013 A comparison of bats and rodents as reservoirs of zoonotic viruses: are bats special? Proc R Soc B 280, 20122753. (doi:10.1098/rspb.2012.2753)

22. Olival KJ, Hosseini PR, Zambrana-Torrelio C, Ross N, Bogich TL, Daszak P. 2017 Host and viral traits predict zoonotic spillover from mammals. Nature 546, 646–650. (doi:10.1038/nature22975)

23. Amman BR et al. 2015 Oral shedding of marburg virus in experimentally infected egyptian fruit bats (rousettus aegyptiacus). J. Wildl. Dis. 51, 113–124. (doi:10.7589/2014-08-198)

24. Middleton DJ, Morrissy CJ, van der Heide BM, Russell GM, Braun MA, Westbury HA, Halpin K, Daniels PW. 2007 Experimental Nipah Virus Infection in Pteropid Bats (Pteropus poliocephalus). J. Comp. Pathol. 136, 266–272. (doi:10.1016/j.jcpa.2007.03.002)

25. Amman BR et al. 2012 Seasonal Pulses of Marburg Virus Circulation in Juvenile Rousettus aegyptiacus Bats Coincide with Periods of Increased Risk of Human Infection. PLOS Pathog. 8, e1002877. (doi:10.1371/journal.ppat.1002877)

26. McFarlane R, Becker N, Field H. 2011 Investigation of the Climatic and Environmental Context of Hendra Virus Spillover Events 1994-2010. PLOS ONE 6, e28374. (doi:10.1371/journal.pone.0028374)

27. Luby SP et al. 2006 Foodborne Transmission of Nipah Virus, Bangladesh. Emerg. Infect. Dis. 12, 1888–1894. (doi:10.3201/eid1212.060732)

28. Pulliam JRC et al. 2012 Agricultural intensification, priming for persistence and the emergence of Nipah virus: a lethal bat-borne zoonosis. J. R. Soc. Interface 9, 89–101. (doi:10.1098/rsif.2011.0223)

29. Glennon EE, Restif O, Sbarbaro SR, Garnier R, Cunningham AA, Suu-Ire RD, Osei-Amponsah R, Wood JLN, Peel AJ. 2018 Domesticated animals as hosts of henipaviruses and filoviruses: A systematic review. Vet. J. 233, 25–34. (doi:10.1016/j.tvjl.2017.12.024)

30. Pourrut X, Kumulungui B, Wittmann T, Moussavou G, Délicat A, Yaba P, Nkoghe D, Gonzalez J-P, Leroy EM. 2005 The natural history of Ebola virus in Africa. Microbes Infect. 7, 1005–1014.

31. Brook CE et al. 2019 Disentangling serology to elucidate henipa- and filovirus transmission in Madagascar fruit bats. J. Anim. Ecol. 0. (doi:10.1111/1365-2656.12985)

32. Pourrut X, Délicat A, Rollin PE, Ksiazek TG, Gonzalez J-P, Leroy EM. 2007 Spatial and Temporal Patterns of Zaire ebolavirus Antibody Prevalence in the Possible Reservoir Bat Species. J. Infect. Dis. 196, S176–S183. (doi:10.1086/520541)

33. Páez DJ, Giles J, McCallum H, Field H, Jordan D, Peel AJ, Plowright RK. 2017 Conditions affecting the timing and magnitude of Hendra virus shedding across pteropodid bat populations in Australia. Epidemiol. Infect., 1–11. (doi:10.1017/S0950268817002138)

34. Drexler JF et al. 2012 Bats host major mammalian paramyxoviruses. Nat. Commun. 3, 796. (doi:10.1038/ncomms1796)

35. Han BA, Schmidt JP, Alexander LW, Bowden SE, Hayman DT, Drake JM. 2016 Undiscovered bat hosts of filoviruses. PLOS Negl Trop Dis 10, e0004815.

36. Schuh AJ et al. 2017 Modelling filovirus maintenance in nature by experimental transmission of Marburg virus between Egyptian rousette bats. Nat. Commun. 8, 14446.

37. Hayman DTS, Bowen RA, Cryan PM, McCracken GF, O’Shea TJ, Peel AJ, Gilbert A, Webb CT, Wood JLN. 2013 Ecology of Zoonotic Infectious Diseases in Bats: Current Knowledge and Future Directions. Zoonoses Public Health 60, 2–21. (doi:10.1111/zph.12000)

38. Sokolow Susanne H. et al. 2019 Ecological interventions to prevent and manage zoonotic pathogen spillover. Philos. Trans. R. Soc. B Biol. Sci. 374, 20180342. (doi:10.1098/rstb.2018.0342)

39. Peel AJ et al. 2013 Use of cross-reactive serological assays for detecting novel pathogens in wildlife: Assessing an appropriate cutoff for henipavirus assays in African bats. J. Virol. Methods 193, 295–303. (doi:10.1016/j.jviromet.2013.06.030)

40. Gilbert AT et al. 2013 Deciphering serology to understand the ecology of infectious diseases in wildlife. Ecohealth 10, 298–313.

41. Wacharapluesadee S, Boongird K, Wanghongsa S, Ratanasetyuth N, Supavonwong P, Saengsen D, Gongal G n., Hemachudha T. 2009 A Longitudinal Study of the Prevalence of Nipah Virus in Pteropus lylei Bats in Thailand: Evidence for Seasonal Preference in Disease Transmission. Vector-Borne Zoonotic Dis. 10, 183–190. (doi:10.1089/vbz.2008.0105)

42. Halpin K et al. 2011 Pteropid bats are confirmed as the reservoir hosts of henipaviruses: a comprehensive experimental study of virus transmission. Am. J. Trop. Med. Hyg. 85, 946–951. (doi:10.4269/ajtmh.2011.10-0567)

43. Wood S. 2006. Generalized additive models: an introduction with R. CRC press.

44. Viechtbauer W. 2010 Conducting meta-analyses in R with the metafor package. J. Stat. Softw. 36, 1–48.

45. Konstantopoulos S. 2011 Fixed effects and variance components estimation in three-level meta-analysis. Res. Synth. Methods 2, 61–76.

46. Nakagawa S, Santos ES. 2012 Methodological issues and advances in biological meta-analysis. Evol. Ecol. 26, 1253–1274.

47. Michonneau F, Brown JW, Winter DJ. 2016 rotl: an R package to interact with the Open Tree of Life data. Methods Ecol. Evol. (doi:10.1111/2041-210X.12593)

48. Paradis E, Claude J, Strimmer K. 2004 APE: analyses of phylogenetics and evolution in R language. Bioinformatics 20, 289–290.

49. Senior AM, Grueber CE, Kamiya T, Lagisz M, O’Dwyer K, Santos ESA, Nakagawa S. 2016 Heterogeneity in ecological and evolutionary meta-analyses: its magnitude and implications. Ecology 97, 3293–3299. (doi:10.1002/ecy.1591)

50. Borenstein M, Hedges LV, Higgins JP, Rothstein HR. 2011 Introduction to meta-analysis. John Wiley & Sons. See http://books.google.com/books?hlen&lr&idJQg9jdrq26wC&oifnd&pgPT14&dqBorenstein+M,+Hedges+L,+Higgins+J,+Rothstein+H+(2005)+Comprehensive+Meta-Analysis&otsVH_-OOjzjB&sigobmoEp98npSeyQvbvsSC_jjdPMc.

51. López-López JA, Marín-Martínez F, Sánchez-Meca J, Noortgate W, Viechtbauer W. 2014 Estimation of the predictive power of the model in mixed-effects meta-regression: A simulation study. Br. J. Math. Stat. Psychol. 67, 30–48.

52. Changula K et al. 2018 Seroprevalence of filovirus infection of Rousettus aegyptiacus bats in Zambia. J. Infect. Dis. 218, S312–S317.

53. Pawęska JT, van Vuren PJ, Kemp A, Storm N, Grobbelaar AA, Wiley MR, Palacios G, Markotter W. 2018 Marburg virus infection in Egyptian rousette bats, South Africa, 2013-2014. Emerg. Infect. Dis. 24, 1134.

54. Plowright RK, Field HE, Smith C, Divljan A, Palmer C, Tabor G, Daszak P, Foley JE. 2008 Reproduction and nutritional stress are risk factors for Hendra virus infection in little red flying foxes (Pteropus scapulatus). Proc. R. Soc. B 275, 861–869. (doi:10.1098/rspb.2007.1260)

55. Washburne AD, Crowley DE, Becker DJ, Olival KJ, Taylor M, Munster VJ, Plowright RK. 2018 Taxonomic patterns in the zoonotic potential of mammalian viruses. PeerJ 6, e5979. (doi:10.7717/peerj.5979)

56. Cressie N, Wikle CK. 2015 Statistics for Spatio-Temporal Data. John Wiley & Sons.

57. Yoccoz NG, Nichols JD, Boulinier T. 2001 Monitoring of biological diversity in space and time. Trends Ecol. Evol. 16, 446–453.

58. Urquhart NS, Kincaid TM. 1999 Designs for detecting trend from repeated surveys of ecological resources. J. Agric. Biol. Environ. Stat., 404–414.

59. Becker DJ, Crowley DE, Washburne AD, Plowright RK. 2019 Data from: Temporal and spatial limitations in global surveillance for bat filoviruses and henipaviruses. (doi:10.5061/dryad.kkwh70s18)

