## Supplemental Material for "Temporal and spatial limitations in global surveillance for bat filoviruses and henipaviruses"

**S1. Systematic search**
**S2. Full reference list**
**S3. Bat phylogeny**
**S4. Temporal patterns in longitudinal sampling and reporting**
**S5. Distribution of sampling duration**
**S6. REM results, $I^2$, and $H^2$**
**S7. Sampling design and reporting practices MEM**
**S8. Spatiotemporal variation in longitudinal studies**

**S1. Systematic search**

Figure S1. The data collection and inclusion process for studies of wild bat filovirus and henipavirus prevalence and seroprevalence (PRISMA diagram). Searches used the following string: (bat* OR Chiroptera*) AND (filovirus OR henipavirus OR "Hendra virus" OR "Nipah virus" OR "Ebola virus" OR "Marburg virus" OR ebolavirus OR marburgvirus). Searches were run during October 2017 and again in August 2019, supplemented by extracting data from references cited in identified studies. Publications were excluded if they did not assess filovirus or henipavirus prevalence or seroprevalence in wild bats or were in languages other than English.

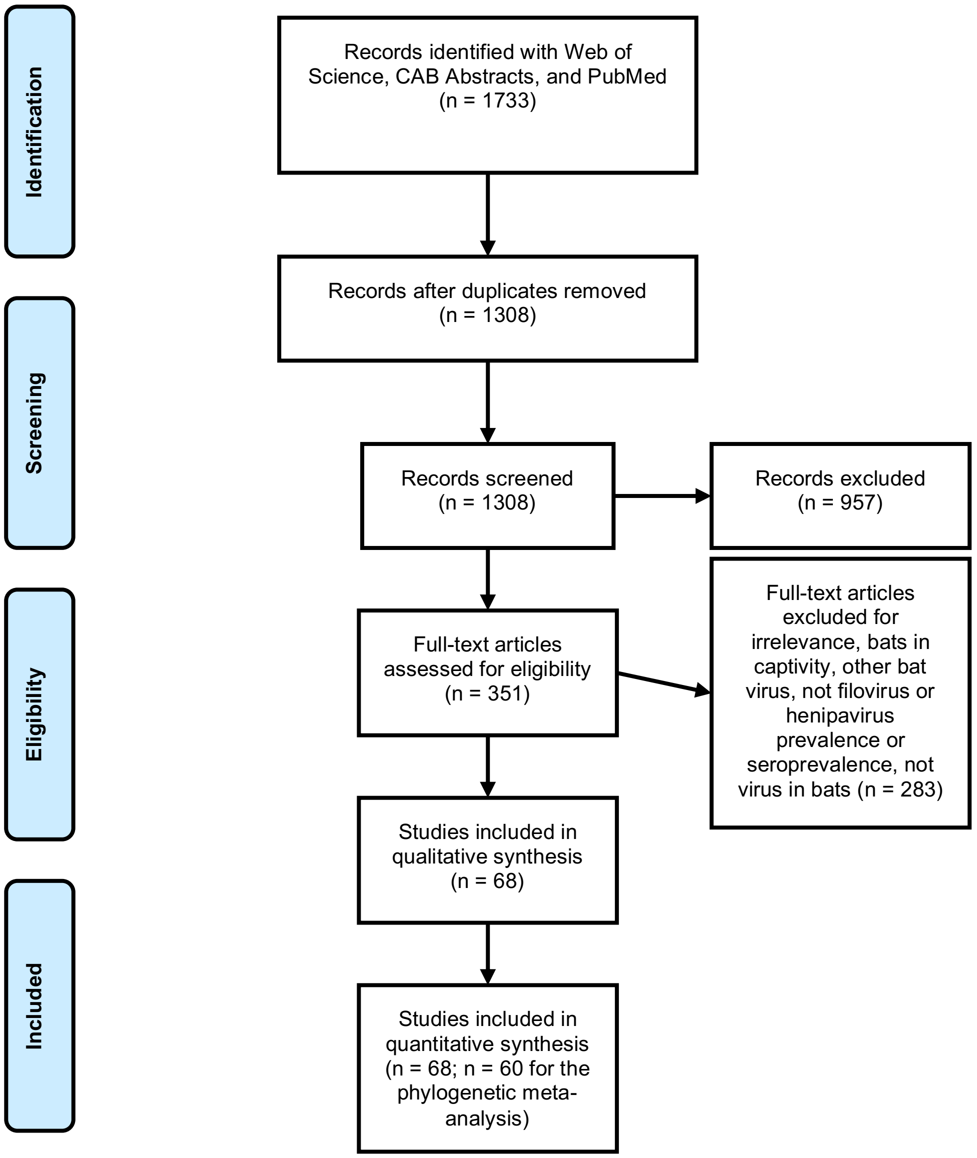

**S2. Full reference list**

1. Amman BR *et al.* 2012 Seasonal Pulses of Marburg Virus Circulation in Juvenile Rousettus aegyptiacus Bats Coincide with Periods of Increased Risk of Human Infection. *PLOS Pathogens* **8**, e1002877. (doi:10.1371/journal.ppat.1002877)

2. Breed AC, Breed MF, Meers J, Field HE. 2011 Evidence of Endemic Hendra Virus Infection in Flying-Foxes (Pteropus conspicillatus)—Implications for Disease Risk Management. *PLOS ONE* **6**, e28816. (doi:10.1371/journal.pone.0028816)

3. Breed AC, Meers J, Sendow I, Bossart KN, Barr JA, Smith I, Wacharapluesadee S, Wang L, Field HE. 2013 The distribution of henipaviruses in Southeast Asia and Australasia: is Wallace's line a barrier to Nipah virus? *PloS one* **8**, e61316.

4. Breed AC, Yu M, Barr JA, Crameri G, Thalmann CM, Wang L-F. 2010 Prevalence of henipavirus and rubulavirus antibodies in pteropid bats, Papua New Guinea. *Emerging infectious diseases* **16**, 1997.

5. Breman JG, Johnson KM, Van Der Groen G, Robbins CB, Szczeniowski MV, Ruti K, Webb PA, Meier F, Heymann DL. 1999 A search for Ebola virus in animals in the Democratic Republic of the Congo and Cameroon: ecologic, virologic, and serologic surveys, 1979–1980. *The Journal of infectious diseases* **179**, S139–S147.

6. Brook CE *et al.* 2019 Disentangling serology to elucidate henipa-and filovirus transmission in Madagascar fruit bats. *Journal of Animal Ecology*

7. Changula K *et al.* 2018 Seroprevalence of filovirus infection of Rousettus aegyptiacus bats in Zambia. *The Journal of infectious diseases* **218**, S312–S317.

8. de Araujo J *et al.* 2017 Antibodies Against Henipa-Like Viruses in Brazilian Bats. *Vector-Borne and Zoonotic Diseases* **17**, 271–274. (doi:10.1089/vbz.2016.2051)

9. De Nys HM *et al.* 2018 Survey of Ebola viruses in frugivorous and insectivorous bats in Guinea, Cameroon, and the Democratic Republic of the Congo, 2015–2017. *Emerging infectious diseases* **24**, 2228.

10. Drexler JF *et al.* 2009 Henipavirus RNA in African bats. *PloS one* **4**, e6367.

11. Drexler JF *et al.* 2012 Bats host major mammalian paramyxoviruses. *Nature communications* **3**, 796.

12. Edson D *et al.* 2019 Time of year, age class and body condition predict Hendra virus infection in Australian black flying foxes (Pteropus alecto). *Epidemiology & Infection* **147**.

13. Edson D *et al.* 2015 Routes of Hendra virus excretion in naturally-infected flying-foxes: implications for viral transmission and spillover risk. *PloS one* **10**, e0140670.

14. Epstein JH, Prakash V, Smith CS, Daszak P, McLaughlin AB, Meehan G, Field HE, Cunningham AA. 2008 Henipavirus infection in fruit bats (Pteropus giganteus), India. *Emerging infectious diseases* **14**, 1309.

15. Epstein JH *et al.* 2006 Feral cats and risk for Nipah virus transmission. *Emerging infectious diseases* **12**, 1178.

16. Field H, de Jong C, Melville D, Smith C, Smith I, Broos A, Kung YHN, McLaughlin A, Zeddeman A. 2011 Hendra virus infection dynamics in Australian fruit bats. *PloS one* **6**, e28678.

17. Forbes KM *et al.* 2019 Bombali Virus in Mops condylurus Bat, Kenya. *Emerging infectious diseases* **25**, 955.

18. Goldspink LK, Edson DW, Vidgen ME, Bingham J, Field HE, Smith CS. 2015 Natural Hendra virus infection in flying-foxes-tissue tropism and risk factors. *PLoS One* **10**, e0128835.

19. Goldstein T *et al.* 2018 The discovery of Bombali virus adds further support for bats as hosts of ebolaviruses. *Nature microbiology* **3**, 1084.

20. Hasebe F *et al.* 2012 Serologic evidence of nipah virus infection in bats, Vietnam. *Emerging infectious diseases* **18**, 536.

21. Hayman DT, Emmerich P, Yu M, Wang L-F, Suu-Ire R, Fooks AR, Cunningham AA, Wood JL. 2010 Long-term survival of an urban fruit bat seropositive for Ebola and Lagos bat viruses. *PloS one* **5**, e11978.

22. Hayman DT, Suu-Ire R, Breed AC, McEachern JA, Wang L, Wood JL, Cunningham AA. 2008 Evidence of henipavirus infection in West African fruit bats. *PloS one* **3**, e2739.

23. Hayman DT, Yu M, Crameri G, Wang L-F, Suu-Ire R, Wood JL, Cunningham AA. 2012 Ebola virus antibodies in fruit bats, Ghana, West Africa. *Emerging infectious diseases* **18**, 1207.

24. He B, Feng Y, Zhang H, Xu L, Yang W, Zhang Y, Li X, Tu C. 2015 Filovirus RNA in fruit bats, China. *Emerging infectious diseases* **21**, 1675.

25. Hsu VP *et al.* 2004 Nipah virus encephalitis reemergence, Bangladesh. *Emerging infectious diseases* **10**, 2082.

26. Iehlé C, Razafitrimo G, Razainirina J. 2007 Henipavirus and Tioman virus antibodies in pteropodid bats, Madagascar. *Emerging infectious diseases* **13**, 159.

27. Jayme SI *et al.* 2015 Molecular evidence of Ebola Reston virus infection in Philippine bats. *Virology journal* **12**, 107.

28. Kajihara M *et al.* 2019 Marburgvirus in Egyptian Fruit Bats, Zambia. *Emerging infectious diseases* **25**, 1577.

29. Karan LS *et al.* 2019 Bombali Virus in Mops condylurus Bats, Guinea. *Emerging infectious diseases* **25**, 1774.

30. Kashiwazaki Y, Na YN, Tanimura N, Imada T. 2004 A solid-phase blocking ELISA for detection of antibodies to Nipah virus. *Journal of virological methods* **121**, 259–261.

31. Kuzmin IV, Niezgoda M, Franka R, Agwanda B, Markotter W, Breiman RF, Shieh W-J, Zaki SR, Rupprecht CE. 2010 Marburg virus in fruit bat, Kenya. *Emerging infectious diseases* **16**, 352.

32. Laing ED *et al.* 2018 Serologic evidence of fruit bat exposure to filoviruses, Singapore, 2011–2016. *Emerging infectious diseases* **24**, 122.

33. Leirs H, Mills JN, Krebs JW, Childs JE, Akaibe D, Woollen N, Ludwig G, Peters CJ, Ksiazek TG. 1999 Search for the Ebola virus reservoir in Kikwit, Democratic Republic of the Congo: reflections on a vertebrate collection. *The Journal of infectious diseases* **179**, S155–S163.

34. Leroy EM *et al.* 2005 Fruit bats as reservoirs of Ebola virus. *Nature* **438**, 575.

35. Li Y *et al.* 2008 Antibodies to Nipah or Nipah-like viruses in bats, China. *Emerging infectious diseases* **14**, 1974.

36. Maganga GD, Bourgarel M, Ebang Ella G, Drexler JF, Gonzalez J, Drosten C, Leroy EM. 2011 Is Marburg virus enzootic in Gabon? *The Journal of infectious diseases* **204**, S800–S803.

37. Negredo A *et al.* 2011 Discovery of an ebolavirus-like filovirus in europe. *PLoS pathogens* **7**, e1002304.

38. Ogawa H *et al.* 2015 Seroepidemiological prevalence of multiple species of filoviruses in fruit bats (Eidolon helvum) migrating in Africa. *The Journal of infectious diseases* **212**, S101–S108.

39. Olival KJ *et al.* 2013 Ebola virus antibodies in fruit bats, Bangladesh. *Emerging infectious diseases* **19**, 270.

40. Paweska JT, van Vuren PJ, Kemp A, Storm N, Grobbelaar AA, Wiley MR, Palacios G, Markotter W. 2018 Marburg virus infection in Egyptian rousette bats, South Africa, 2013–2014. *Emerging infectious diseases* **24**, 1134.

41. Peel AJ *et al.* 2012 Henipavirus neutralising antibodies in an isolated island population of African fruit bats. *PloS one* **7**, e30346.

42. Peel AJ *et al.* 2013 Continent-wide panmixia of an African fruit bat facilitates transmission of potentially zoonotic viruses. *Nature communications* **4**, 2770.

43. Plowright RK, Field HE, Smith C, Divljan A, Palmer C, Tabor G, Daszak P, Foley JE. 2008 Reproduction and nutritional stress are risk factors for Hendra virus infection in little red flying foxes (Pteropus scapulatus). *Proceedings of the Royal Society B: Biological Sciences* **275**, 861–869.

44. Pourrut X, Delicat A, Rollin PE, Ksiazek TG, Gonzalez J-P, Leroy EM. 2007 Spatial and temporal patterns of Zaire ebolavirus antibody prevalence in the possible reservoir bat species. *The Journal of infectious diseases* **196**, S176–S183.

45. Pourrut X, Souris M, Towner JS, Rollin PE, Nichol ST, Gonzalez J-P, Leroy E. 2009 Large serological survey showing cocirculation of Ebola and Marburg viruses in Gabonese bat populations, and a high seroprevalence of both viruses in Rousettus aegyptiacus. *BMC infectious diseases* **9**, 159.

46. Pulliam JR *et al.* 2011 Agricultural intensification, priming for persistence and the emergence of Nipah virus: a lethal bat-borne zoonosis. *Journal of the Royal Society Interface* **9**, 89–101.

47. Rahman SA *et al.* 2013 Risk factors for Nipah virus infection among pteropid bats, Peninsular Malaysia. *Emerging infectious diseases* **19**, 51.

48. Ramírez de Arellano E *et al.* 2019 First Evidence of Antibodies Against Lloviu Virus in Schreiber's Bent-Winged Insectivorous Bats Demonstrate a Wide Circulation of the Virus in Spain. *Viruses* **11**, 360.

49. Reynes J-M *et al.* 2005 Nipah virus in Lyle's flying foxes, Cambodia. *Emerging infectious diseases* **11**, 1042.

50. Saéz AM *et al.* 2015 Investigating the zoonotic origin of the West African Ebola epidemic. *EMBO molecular medicine* **7**, 17–23.

51. Sendow I, Field HE, Adjid A, Ratnawati A, Breed AC, Morrissy C, Daniels P. 2010 Screening for Nipah virus infection in West Kalimantan province, Indonesia. *Zoonoses and public health* **57**, 499–503.

52. Sendow I, Field HE, Curran J. 2006 Henipavirus in Pteropus vampyrus bats, Indonesia. *Emerging infectious diseases* **12**, 711.

53. Sendow I, Ratnawati A, Taylor T, Adjid RA, Saepulloh M, Barr J, Wong F, Daniels P, Field H. 2013 Nipah virus in the fruit bat Pteropus vampyrus in Sumatera, Indonesia. *PLoS One* **8**, e69544.

54. Shirai J, Sohayati AL, ALI ALM, Suriani MN, Taniguchi T, Sharifah SH. 2007 Nipah virus survey of flying foxes in Malaysia. *Japan Agricultural Research Quarterly: JARQ* **41**, 69–78.

55. Swanepoel R *et al.* 2007 Studies of reservoir hosts for Marburg virus. *Emerging infectious diseases* **13**, 1847.

56. Taniguchi S *et al.* 2011 Reston Ebolavirus antibodies in bats, the Philippines. *Emerging infectious diseases* **17**, 1559.

57. Towner JS *et al.* 2009 Isolation of genetically diverse Marburg viruses from Egyptian fruit bats. *PLoS pathogens* **5**, e1000536.

58. Towner JS *et al.* 2007 Marburg virus infection detected in a common African bat. *PloS one* **2**, e764.

59. Wacharapluesadee S, Boongird K, Wanghongsa S, Ratanasetyuth N, Supavonwong P, Saengsen D, Gongal GN, Hemachudha T. 2010 A longitudinal study of the prevalence of Nipah virus in Pteropus lylei bats in Thailand: evidence for seasonal preference in disease transmission. *Vector-Borne and Zoonotic Diseases* **10**, 183–190.

60. Wacharapluesadee S *et al.* 2005 Bat Nipah virus, Thailand. *Emerging infectious diseases* **11**, 1949.

61. Wacharapluesadee S *et al.* 2015 Surveillance for Ebola virus in wildlife, Thailand. *Emerging infectious diseases* **21**, 2271.

62. Wacharapluesadee S, Samseeneam P, Phermpool M, Kaewpom T, Rodpan A, Maneeorn P, Srongmongkol P, Kanchanasaka B, Hemachudha T. 2016 Molecular characterization of Nipah virus from Pteropus hypomelanus in Southern Thailand. *Virology journal* **13**, 53.

63. Yadav PD, Raut CG, Shete AM, Mishra AC, Towner JS, Nichol ST, Mourya DT. 2012 Detection of Nipah virus RNA in fruit bat (Pteropus giganteus) from India. *The American journal of tropical medicine and hygiene* **87**, 576–578.

64. Yadav PD *et al.* 2019 Nipah Virus Sequences from Humans and Bats during Nipah Outbreak, Kerala, India, 2018. *Emerging infectious diseases* **25**, 1003.

65. Yang X-L *et al.* 2017 Genetically diverse filoviruses in Rousettus and Eonycteris spp. bats, China, 2009 and 2015. *Emerging infectious diseases* **23**, 482.

66. Yob JM *et al.* 2001 Nipah virus infection in bats (order Chiroptera) in peninsular Malaysia. *Emerging infectious diseases* **7**, 439.

67. Young PL, Halpin K, Selleck PW, Field H, Gravel JL, Kelly MA, Mackenzie JS. 1996 Serologic evidence for the presence in Pteropus bats of a paramyxovirus related to equine morbillivirus. *Emerging infectious diseases* **2**, 239.

68. Yuan J, Zhang Y, Li J, Zhang Y, Wang L-F, Shi Z. 2012 Serological evidence of ebolavirus infection in bats, China. *Virology journal* **9**, 236.

### S3. Bat phylogeny

Figure S2. Phylogeny of the 215 bat species included in the phylogenetic meta-analysis of filovirus and henipavirus prevalence and seroprevalence.

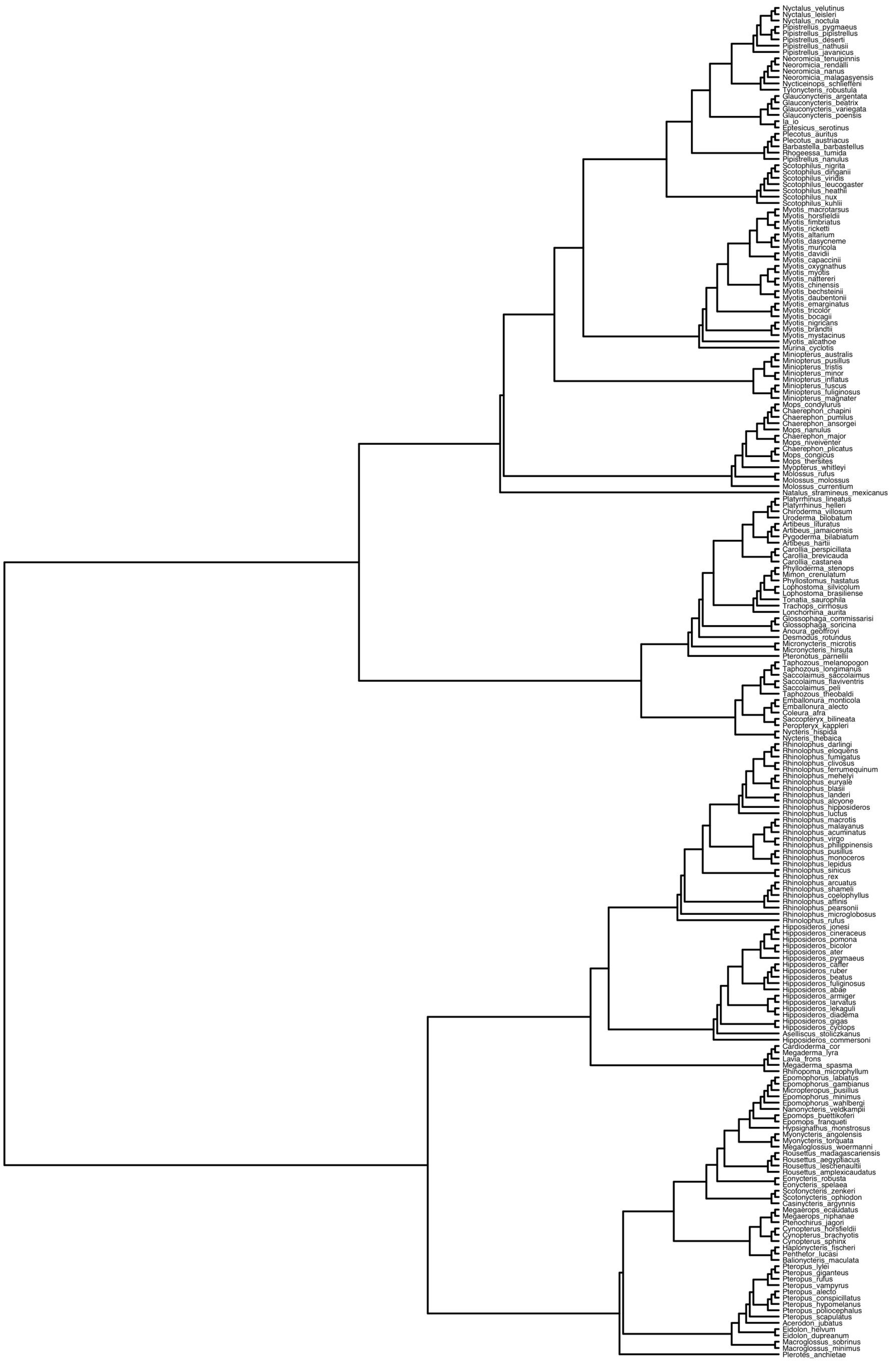

**S4. Temporal patterns in longitudinal sampling and reporting**

Figure S3. Relationship between publication year and the proportion of studies reporting longitudinal data. Fitted values (line) and 95% confidence intervals (grey) from a generalized additive model (binomial errors) are shown with data scaled by the number of studies per year.

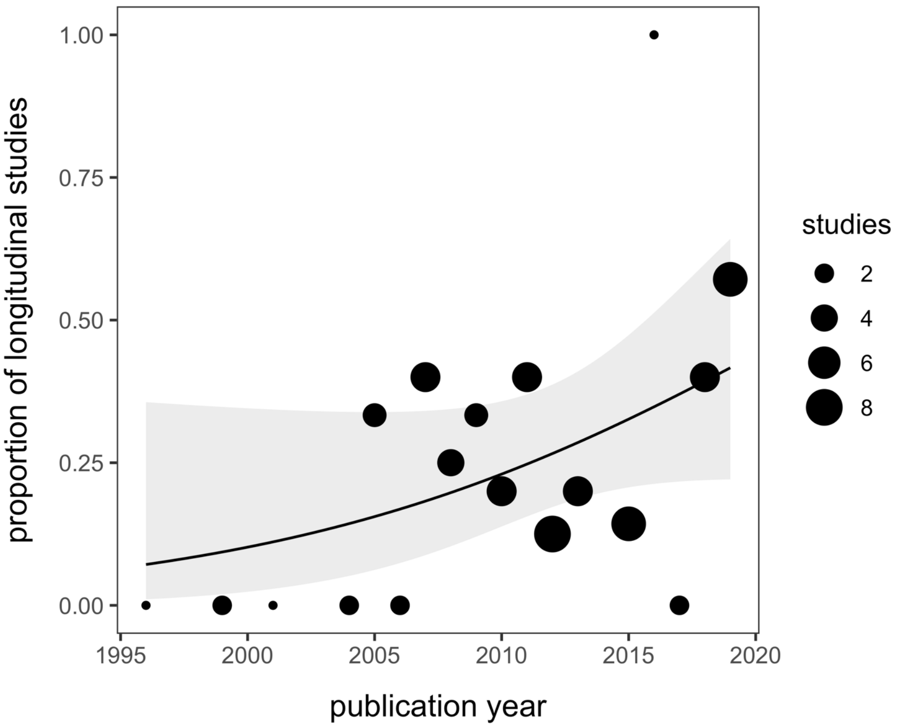

**S5. Distribution of sampling duration**

Figure S4. Distribution of study duration in years for longitudinal studies; the mean (2.5) is shown with the black vertical line.

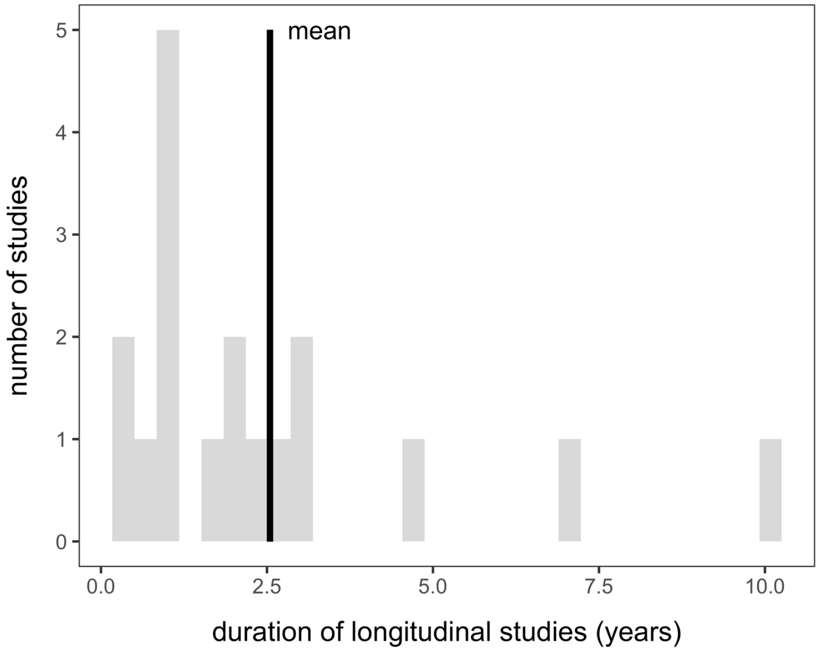


Figure S5. Distribution of days represented by each pooled viral detection estimate; the mean (644) is shown with the black vertical line.

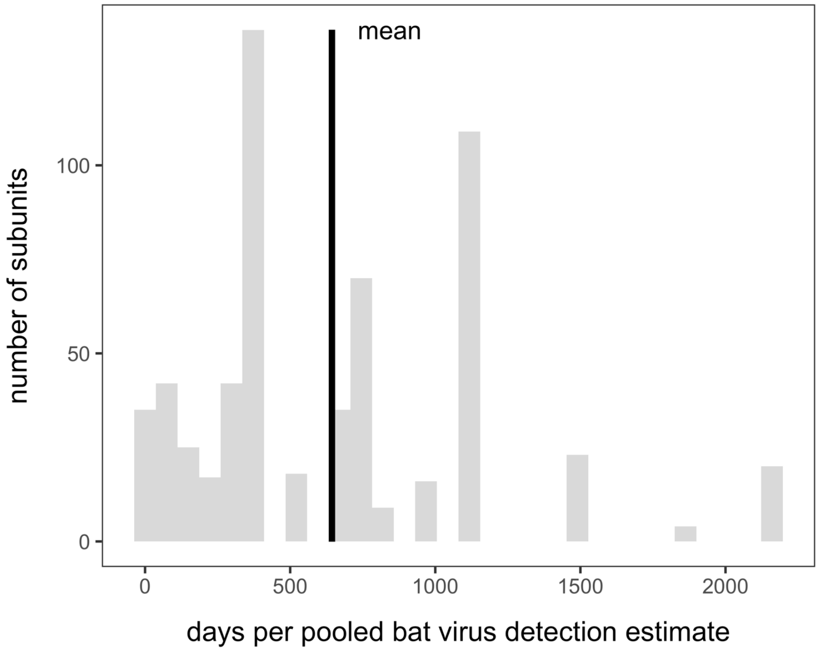

### S6. REM results, $I^2$, and $H^2$

Table S1. Results from the REM fit to all data and each data subset, using the random effects outlined in the main text. We show Cochran's $Q$, the associated $p$ value, estimates of total $I^2$ and the proportion of unknown variation not attributable to sampling variance for each random effect ($I^2_{species}$, $I^2_{study}$, and $I^2_{subunit}$), and phylogenetic heritability ($H^2$).

| **Data** | $Q$ | $p$ | $I^2$ | $I^2_{species}$ | $I^2_{study}$ | $I^2_{subunit}$ | $H^2$ |
|---|---|---|---|---|---|---|---|
| All data | 6929 | <0.001 | 0.91 | 0.41 | 0.34 | 0.15 | 0.45 |
| Filovirus prevalence | 340 | <0.001 | 0.53 | 0.03 | 0.22 | 0.27 | 0.07 |
| Filovirus seroprevalence | 2337 | <0.001 | 0.85 | 0 | 0.66 | 0.2 | 0 |
| Henipavirus prevalence | 401 | <0.001 | 0.63 | 0.30 | 0.16 | 0.16 | 0.48 |
| Henipavirus seroprevalence | 1926 | <0.001 | 0.96 | 0.66 | 0.17 | 0.14 | 0.68 |

**S7. Sampling design and reporting practices MEM**

Table S2. ANOVA table from the MEM with a three-way interaction between sampling design and reporting practices, virus taxa, and detection method. For each term in the model, we provide Cochran's *Q*, the associated degrees of freedom, and the *p* value.

| **Model term** | ***Q*** | ***df*** | ***p*** |
|---|---|---|---|
| Sampling and reporting practices | 0.20 | 2 | 0.91 |
| Virus taxa | 0.09 | 1 | 0.77 |
| Detection method | 5.41 | 1 | 0.02 |
| Sampling practices * virus taxa | 0.46 | 2 | 0.79 |
| Sampling practices * detection method | 3.84 | 2 | 0.15 |
| Virus taxa * detection method | 0.03 | 1 | 0.87 |
| Sampling practices * virus taxa * detection method | 5.36 | 2 | 0.07 |

Table S3. Post-hoc analysis for the marginally significant interaction between sampling design and reporting practices, detection method, and virus taxa. MEMs with the same random effects were fit to each data subset (filovirus prevalence=193, filovirus seroprevalence=302, henipavirus prevalence=244, henipavirus seroprevalence=280). For each MEM, we provide the *Q* statistic and *p* value for sampling design and reporting practices.

| **Data subset** | $Q_2$ | ***p*** |
|---|---|---|
| Filovirus prevalence | 0.12 | 0.94 |
| Filovirus seroprevalence | 10.30 | 0.006 |
| Henipavirus prevalence | 2.03 | 0.36 |
| Henipavirus seroprevalence | 1.35 | 0.51 |

**S8. Spatiotemporal variation in longitudinal studies**

Table S4. Signal of spatial variation within longitudinal studies. MEMs with the same random effects were fit to each data subset (filovirus prevalence=22, filovirus seroprevalence=73, henipavirus prevalence=88, henipavirus seroprevalence=90) with sampling location as a predictor. For each model, we provide the omnibus $Q$ statistic and $p$ value.

| **Data subset** | ***Q*** | ***df*** | ***p*** |
|---|---|---|---|
| Filovirus prevalence | 0.03 | 2 | 0.99 |
| Filovirus seroprevalence | 42.39 | 7 | <0.001 |
| Henipavirus prevalence | 23.96 | 10 | 0.008 |
| Henipavirus seroprevalence | 22.39 | 9 | 0.008 |

Table S5. Signal of temporal variation within longitudinal studies. MEMs with the same random effects were fit to each data subset (filovirus prevalence=22, filovirus seroprevalence=73, henipavirus prevalence=88, henipavirus seroprevalence=90) with month as a categorical predictor. For each model, we provide the omnibus $Q$ statistic and $p$ value.

| **Data subset** | ***Q*** | ***df*** | ***p*** |
|---|---|---|---|
| Filovirus prevalence | 1.44 | 6 | 0.96 |
| Filovirus seroprevalence | 23.42 | 11 | 0.02 |
| Henipavirus prevalence | 52.35 | 11 | <0.001 |
| Henipavirus seroprevalence | 13.12 | 11 | 0.29 |
